# Biophysical and molecular mechanisms responsible for phytoplankton sinking in response to starvation

**DOI:** 10.1101/2025.05.04.652135

**Authors:** Yanqi Wu, Vieyiti K. Kouadio, Thomas R. Usherwood, Justin Li, Margaret Bisher, Reshum Aurora, Aaron Z. Lam, Alice R. Lam, Abigail K. R. Lytton-Jean, Scott R. Manalis, Teemu P. Miettinen

## Abstract

Marine phytoplankton face eco-evolutionary pressure to regulate their vertical position in the ocean to access light, which is abundant towards the surface, and nutrients, which are found deeper down the water column. All phytoplankton experience gravitational sinking, which can contribute to their vertical migration. However, the biophysical and molecular mechanisms that impact gravitational sinking have not been systematically characterized across taxa and environmental conditions. Here, we combine simulations with measurements of cell mass, volume, and composition to investigate the effects of nutrient availability on gravitational sinking in 9 representative unicellular pico- and nanoplankton species. We find that gravitational sinking becomes faster in most species when starved, but the biophysical changes responsible for this vary across species and starvation conditions. For example, the faster sinking of *Chaetoceros calcitrans* is nearly exclusively driven by cell density whereas that of *Emiliania huxleyi* is due to cell volume. On the molecular level, the altered sinking is predominantly attributed to changes in cellular dry contents, rather than water. For example, starch accumulation increases sinking in 3 green algae species, and lipid accumulation decreases sinking in *Phaeodactylum tricornutum*. Overall, our work reveals that phytoplankton physiology has evolved multiple mechanisms that impact gravitational sinking in response to starvation, possibly to support the vertical migration of the cell.

## INTRODUCTION

Phytoplankton are key primary producers in the oceans that support marine food webs and drive carbon fixation ^1,2^. Their growth and fitness depend on photosynthesis-derived energy and seawater nutrients– resources that are unevenly distributed in the ocean: light is more abundant towards the surface, while nutrients are more concentrated deeper in the water column ^3,4^. This generates eco-evolutionary pressure for phytoplankton to migrate vertically in the water column to meet their energy and nutrient requirements ^5^. Nutrient-limited phytoplankton may sink deeper in the water column in search of nutrients, simultaneously contributing to the downward flux of organic carbon. Although it is unclear if phytoplankton regulate their sinking to reach more nutrients, thus achieving a fitness advantage, or if changes in cell sinking are simply byproducts of other metabolic changes, the vertical movement of cells in the ocean has ecological consequences. The vertical movement of phytoplankton is predicted to have a significant impact on primary production and nutrient cycles in the oceans ^2,6–8^, and field observations have confirmed the sinking of single cells and small particles (<100 µm cell aggregates) as contributors to ocean carbon fluxes ^9–12^. Yet, the degree to which phytoplankton sinking is impacted by nutrient limitations, as well as the mechanisms responsible for such phenotypic response, have not been systematically characterized across taxa (Fig 1A).

**Figure 1.**
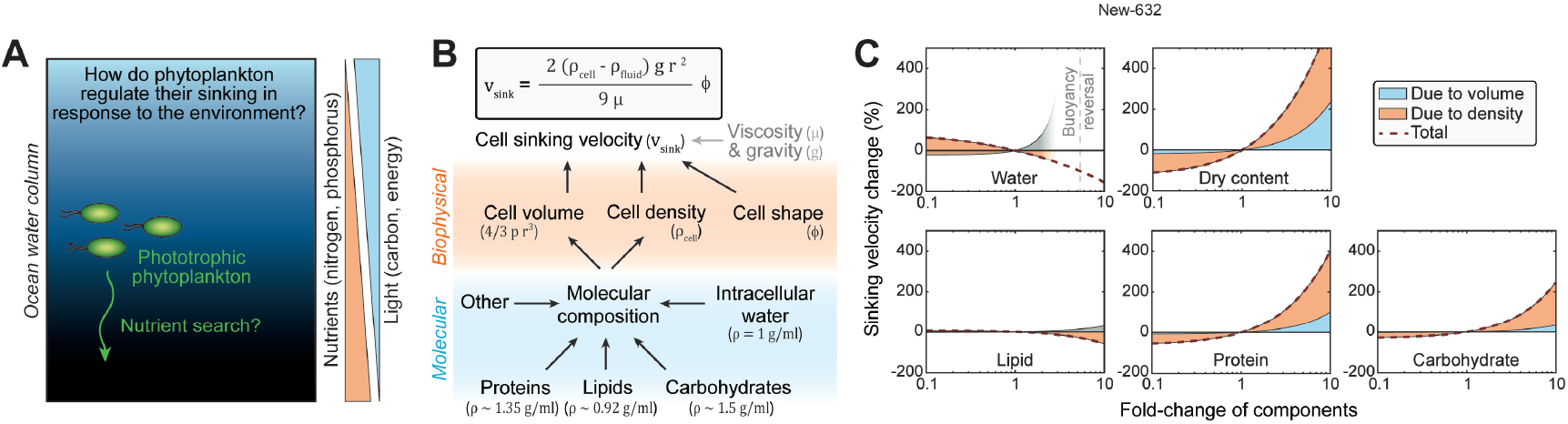
Gravitational sinking velocity can be simulated using a simple model of cell composition. **(A)** Schematic of study question. **(B)** A simple physical model linking cell composition to cell sinking. Gravitational sinking velocity is determined by Stokes’ law (top). On biophysical level, cell sinking is dependent of cell volume, density, and shape (middle). On molecular level (bottom), cell volume and density are primarily dependent on cellular water, protein, lipid, and carbohydrate content. **(C)** Simulations of cell sinking velocity in a representative diatom (*Phaeodactylum tricornutum*) as a function of cell composition. Sinking velocity changes due to cell volume and cell density are separated in blue and orange, respectively. Changes in dry content refer to corresponding changes in all other contents except water.

Here, we focus on gravitational sinking as a potential mechanism for vertical migration. While motile plankton can achieve faster migration with phototaxis than with gravitational sinking, gravitational sinking is experienced by all species and may act as an energy-efficient migration mechanism. The gravitational sinking velocity (*v*_*sink*_, from here on referred to simply as sinking) of a cell can be derived from Stokes’ law ^13–15^,

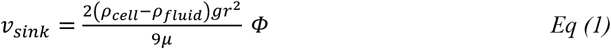

where *ρ*_*cell*_ is the density of the cell, *ρ*_*fluid*_ is the density of the surrounding fluid, *g* is the gravitational acceleration, *r* is the equivalent spherical radius of the cell, μ is the dynamic viscosity of the fluid, and *Φ* is the correction factor for non-spherical shape ^14,16^. Biophysical properties of a cell, namely cell volume, density, and shape, define sinking (Fig 1B), and could determine the sinking of cell aggregates similarly. At the molecular level, we model a cell as the collection of all its intracellular molecules, where the cell volume is the sum volume of individual molecules, and the cell density is the weighted average of the densities of those molecules (Fig 1B). These molecules can be separated into water and dry contents of the cell. For most phytoplankton, especially those without large silica and calcium carbonate shells/walls, most cellular dry contents fall into 3 groups –proteins, lipids, and carbohydrates, each with distinct densities (proteins ∼ 1.35 g/mL, lipids ∼ 0.92 g/mL, carbohydrates ∼ 1.5 g/mL, Table S1) ^17,18^. Previous works have used similar models of cell composition ^19^, and here it provides a framework that relates cell’s molecular content to its biophysical properties and sinking (Fig 1B).

Studies of cell sinking using sedimentation columns or timelapse imaging have shown that many species alter their sinking in response to environmental conditions, such as starvation ^20–25^. However, the mechanisms responsible for these sinking changes are, in most cases, unknown. Many phytoplankton have also been shown to change their molecular composition in response to starvation ^26–30^. Such compositional changes could alter cell sinking, as also suggested by modeling studies ^19^, but there is limited experimental evidence that quantitatively links changes in cell size and composition to cell sinking. Notable exceptions to this are the species *Pyrocystis noctiluca*, which can undergo long vertical migration due to water content regulation ^31^, and *Tetraselmis sp.* which can sink faster when starved due to decreased cellular water content and increased carbohydrate content ^32^.

Here, we study how gravitational sinking responds to different nutrient limitations across a range of motile and non-motile unicellular eukaryotic marine phytoplankton. Using simulations and experiments, we connect cell sinking to the regulation of cell’s biophysical properties and molecular composition. Our work reveals multiple mechanisms responsible for starvation-induced sinking in different phytoplankton species, suggesting that phytoplankton may have evolved several solutions to support their vertical migration.

## RESULTS

### Simulation of phytoplankton sinking velocity

To understand how cell sinking could be impacted by molecular composition, we simulated sinking for cells with different compositions (Fig 1B). Our simulations relied on the previously reported macromolecular composition and size of a typical green algae (*Dunaliella tertiolecta*, Fig S1A, Table S2) or a typical diatom (*Chaetoceros calcitrans* or *Phaeodactylum tricornutum*, Figs 1C, S1B). We then systematically varied the amount of each major intracellular component (lipid, carbohydrate, protein, and water) while keeping other components constant. In addition, we varied all components except water simultaneously in order to examine the effect of dry contents on cell sinking. For each simulated variation, we determined the fraction of the sinking change caused by cell volume and cell density using a first order Taylor linear approximation. The results revealed that: i) faster sinking is more readily achieved by gaining cellular dry contents, specifically carbohydrates and proteins, rather than by decreasing water or lipid content, ii) change in any single cellular component will alter sinking more due to changes in cell density than changes in cell volume, and iii) the reversal of cell buoyancy (decreasing cell density below that of seawater) requires dramatic molecular changes such as large accumulation of water (>4-fold) or lipids (>10-fold) without additional protein or carbohydrate accumulation. These conclusions were not sensitive to the cell composition differences observed between species (Fig S1). We note that our model groups the rest of cellular contents as ‘other’ and, due to the complex nature of this ‘other’ group, we assume it to be constant in density and volume.

### Phytoplankton increase sinking velocity in response to starvation

We have previously established an approach for determining the sinking of pico- and nanoplankton species by applying Stokes’ law to single-cell measurements of cell mass and volume ^32–34^. To examine starvation-induced changes in cell sinking across the tree of life, we selected 9 phototrophic unicellular eukaryotic marine pico- and nanoplankton species and we cultured the cells under photoautotrophic conditions in both high and low nutrient (*i.e.*, starvation) media (Fig 2A, Table S3). Following a 5-day culture, we measured single-cell volumes using Coulter counter, single-cell buoyant masses using a suspended microchannel resonator (SMR), and cell shapes using microscopy. Sinking velocities of a cell population were inferred from population averages of hundreds of single cell measurements (Figs S2A,B). We also confirmed that low nutrient condition resulted in starvation (Fig S3A), and that cell shapes did not change between conditions (Fig S3B). Overall, our analysis focused on species which did not display extensive aggregation, and our data reflects single cells that are either proliferating (high nutrient condition) or starving (low nutrient condition).

**Figure 2.**
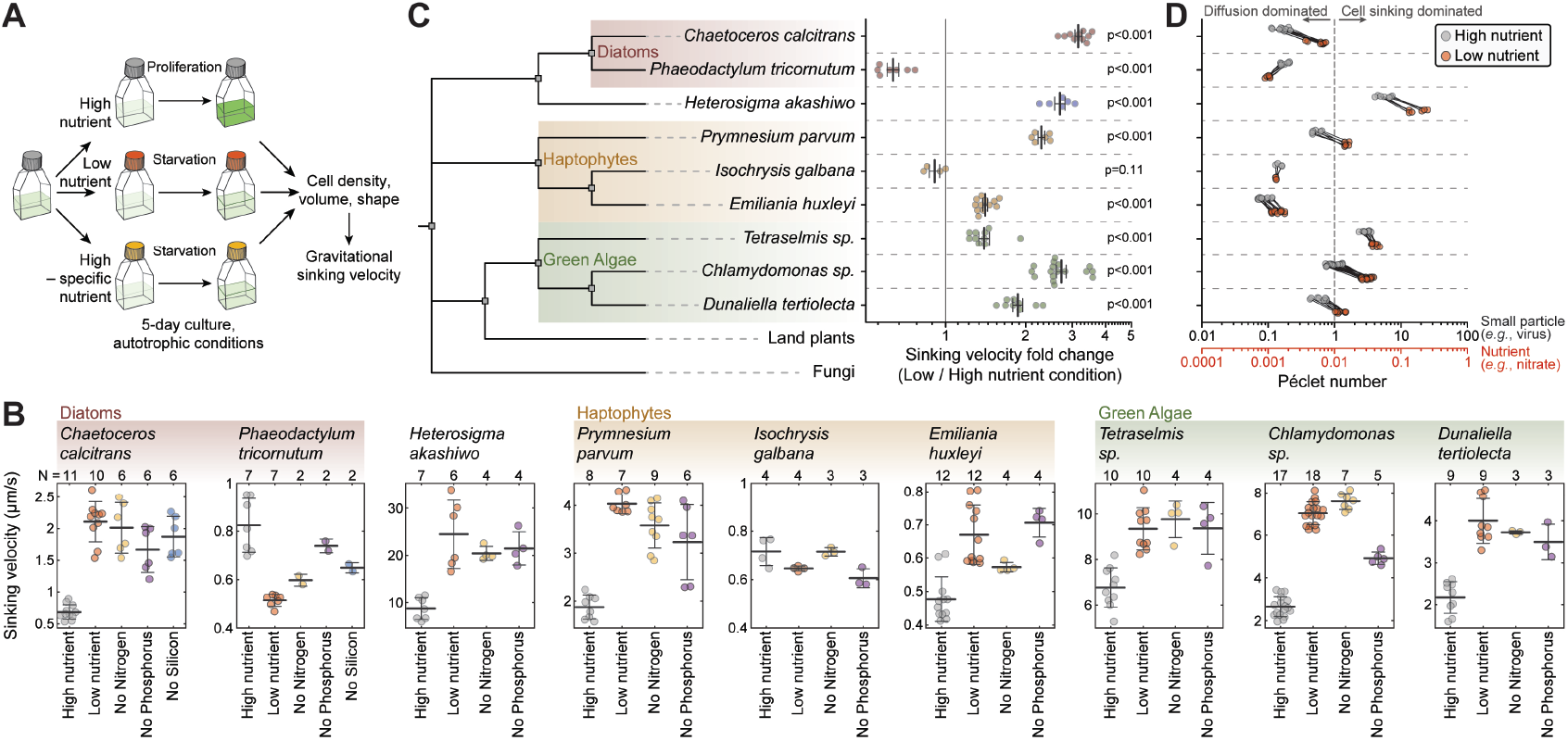
Cell sinking increases in response to starvation across many genetically distinct, unicellular marine phytoplankton. **(A)** Schematic of experimental setup. **(B)** Gravitational sinking velocity in indicated species following 5-day culture under indicated nutrient conditions. N depicts the number of independent cultures (dots), bar and whiskers depict mean±SD. **(C)** Phylogenetic tree of the studied species and the relative sinking velocity change between low (starving) and high (proliferating) nutrient conditions in each species. Land plants and fungi are show for reference. Same data as in (B), dots depict separate cultures, bar and whiskers depict mean±SEM, p-value obtained by t-test comparison to the value of 1. **(D)** Péclet numbers for high (grey dots) and low nutrient (orange dots) conditions, as calculated for the diffusion of nitrate (red x-axis) or a virus particle (black x-axis). Species are in the same order as in (C). Péclet number > 1 indicates that the encounter rate of the cell and the particle (nitrate or virus) is dominated by cell sinking rather than particle diffusion.

Most tested phytoplankton species (7 out of 9) displayed increased sinking when starved for nutrients (Figs 2B, C). For example, in the diatom *Chaetoceros calcitrans*, the toxic raphidophyte *Heterosigma akashiwo*, and the green alga *Chlamydomonas sp.,* sinking increased approximately 3-fold under low nutrient conditions compared to high nutrient conditions. In *Heterosigma akashiwo*, the largest species studied, sinking reached up to ∼25 µm/s when starved. We also observed that one diatom species, *Phaeodactylum tricornutum*, that displayed decreased sinking upon starvation. Only the haptophyte *Isochrysis galbana* did not change sinking significantly following starvation (Figs 2C), and this species was excluded from future analyses. Overall, our results indicate that most marine phytoplankton alter their sinking in response to a general nutrient starvation, but the effect magnitude is species-specific even within phylogenetic clades.

### Starvation increases phytoplankton Péclet numbers, promoting small particle encounters

In environments where nutrients are sparse, nutrient uptake is limited by the rate of diffusion. However, cells can promote their nutrient acquisition by movement. To evaluate if the observed sinking changes are sufficient to promote nutrient acquisition, we calculated the Péclet number (Pe) for a representative small nutrient (nitrate). Pe represents the relative contribution of diffusion and cell sinking to mass transport, with Pe << 1 indicating that diffusion-dominated transport, and Pe >> 1 indicating sinking-dominated transport. Although Pe increased up to 4-fold under low nutrient conditions, the Pe for nitrate remained very small (Fig 2D), indicating that cell sinking does not meaningfully increase nutrient acquisition in the species studied here. We note that this only reflects short timescales when the local nutrient environment remains constant, and over long timescales cells can sink into deeper, nutrient richer waters. On the other hand, particles, such as viruses or other cells, diffuse significantly more slowly than nutrients, leading to Pe_virus_ ≥ 1, for many species when starved (Fig 2D). For example, the toxic haptophyte *Prymnesium parvum* shifted from a diffusion-dominated regime of viral encounters (Pe_virus_ = 0.53±0.03, mean±SEM) to convection-dominated regime of viral interactions (Pe_virus_ = 1.50±0.04, mean±SEM) when starved. Thus, the sinking we observe may increase cell encounters with particles, such as viruses, other cells, or marine snow.

### Sinking velocity is responsive to multiple nutrients

Next, we examined whether the starvation-induced changes in cell sinking were specific to limitation of a single nutrient. More specifically, we used high nutrient media that lacks either nitrogen, phosphorus, or silicon (only for diatoms), and we verified that these conditions result in decreased proliferation (Fig S3A). For all species, we observed qualitatively similar results between starvation by overall low nutrient level and by specific nutrients (Fig 2B). However, in some species, the magnitude of sinking change varied between specific nutrient starvations. For example, the sinking change in *Chaetoceros calcitrans* was greater when starved for nitrogen (p=0.016, paired t-test) or silicon (p=0.003, paired t-test) than when starved for phosphorus. In contrast, in *Emiliania huxleyi*, sinking increased more when starved for phosphorus than nitrogen (p=0.005, paired t-test). Overall, cell sinking is responsive to multiple nutrients, although in some species the effect size varies between different limiting nutrients.

To examine whether the observed changes in sinking were caused by cell death, which can increase cell density ^33^, we resupplied nutrients to the starved cultures in a subset of species studied. The results revealed that sinking could be rescued with nutrient resupply (Fig S4). This is consistent with previous work examining cell size and proliferation recovery following starvation in *Tetraselmis sp.* and *Emiliania huxleyi* ^32,35^.

### Biophysical mechanisms for sinking velocity changes are species-specific

Cell sinking is defined by cell volume, density, and shape, the last of which did not change between conditions (Fig S3B). Here, we consider cell volume as an indicator of total cellular contents, cell density as an indicator of cell composition (Fig 1B), and we examine how sinking is impacted by cell volume and density. We overlaid the measured cell volume and density changes with the resultant sinking (Fig 3A). This revealed that cell density increased in nearly all species upon starvation, except *Emiliania huxleyi*, which displayed a constant density, and *Phaeodactylum tricornutum*, which decreased cell density. Many phytoplankton species also increased their cell volume upon starvation, as has been previously reported for *Emiliania huxleyi* ^35^.

**Figure 3.**
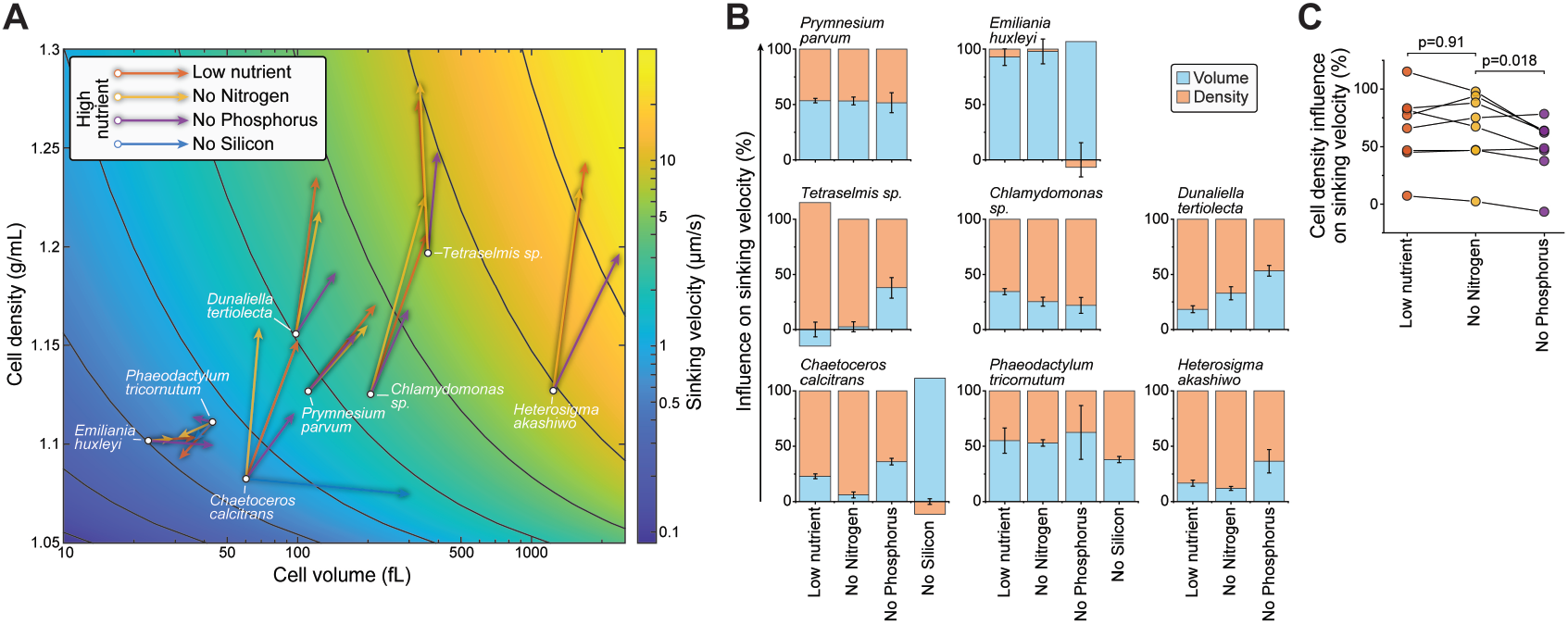
Biophysical changes responsible for starvation-induced cell sinking. **(A)** Changes in cell density and cell volume when comparing high nutrient condition (base of each arrow) to indicated starvation conditions (tip of each arrow). Gravitational sinking velocities are indicated in color-coded background. **(B)** The relative influence of cell volume (blue) and cell density (orange) changes on cell sinking velocity changes under each starvation condition. Negative values imply that volume and density changes have opposite effects on the sinking velocity. Data depicts mean±SEM. **(C)** The relative influence of cell density changes on cell sinking velocity changes under indicated starvation conditions. Dots depict different species, data from same species are connected by a line. p-value obtained by Student’s t-test (N = 8 species). Low nutrient condition and nitrogen starvation appear similar, most likely because nitrogen is the limiting nutrient for most species under the low nutrient conditions.

We then compared how cell volume and density changes influence sinking under each starvation condition. This revealed that green algae, i.e., *Tetraselmis sp*., *Chlamydomonas sp*., and *Dunaliella tertiolecta*, as well as *Heterosigma akashiwo*, relied predominantly on cell density regulation to adjust their sinking (Fig 3B). In contrast, *Phaeodactylum tricornutum* and *Prymnesium parvum* relied approximately evenly on cell volume and density regulation to adjust sinking, whereas *Emiliania huxleyi* relied exclusively on cell volume regulation to adjust sinking. Thus, both cell volume and density regulation can function as the biophysical basis for adjusting sinking, depending on the species.

Starvation for different nutrients resulted in different cell volume and density responses. Most notably, in *Chaetoceros calcitrans*, nitrogen starvation increased cell density with little effect on cell volume, whereas silicon starvation increased cell volume rather than cell density (Fig 3A-B). A broader comparison across all species revealed that nitrogen starvation was more prone to alter cell sinking due to cell density, when compared to phosphorus starvation (Fig 3C).

### Increased cell sinking is driven by dry content accumulation

In theory, the observed changes in sinking, as well as biophysical properties, must be attributed to changes in specific cellular molecules. Our simulations of cell sinking suggested that an overall accumulation of cell dry contents can increase sinking more effectively than the loss of water or lipids (Fig 1C). Motivated by this, we sought to examine if the starvation-driven increases in sinking were driven by dry content accumulation or by loss of intracellular water. Using a previously established approach ^32^, we measured cellular water and dry contents. Most species increased their water and dry content following starvation, except for low nutrient starved *Chaetoceros calcitrans* and *Tetraselmis sp*., and nitrogen starved *Chlamydomonas sp.* (Fig S5A). These results suggest a separate regulation of phytoplankton’s water and dry contents, as observed in other model systems ^36^. We note that *Heterosigma akashiwo* was excluded from this and future experiments due to technical reasons.

We then compared how cellular water and dry content changes influence cell sinking. This revealed that nearly all starvation-driven sinking increases were caused by increases in cellular dry contents (Fig 4). In contrast, cellular water content, which increased in most species following starvation, had a negative influence on sinking (Fig 4). Only in low nutrient starved *Tetraselmis sp.* did the loss of cellular water contribute positively and significantly to the sinking increase (p=0.024, one sample t-test). Therefore, while starvation induced species-and condition-specific changes to cellular water content, cellular water content was rarely an important contributor to sinking. Instead, the increases in sinking were nearly exclusively driven by changes in cellular dry contents. For a breakdown of cellular dry content into its volume and density, and their separate impact on sinking, see Fig S5.

**Figure 4.**
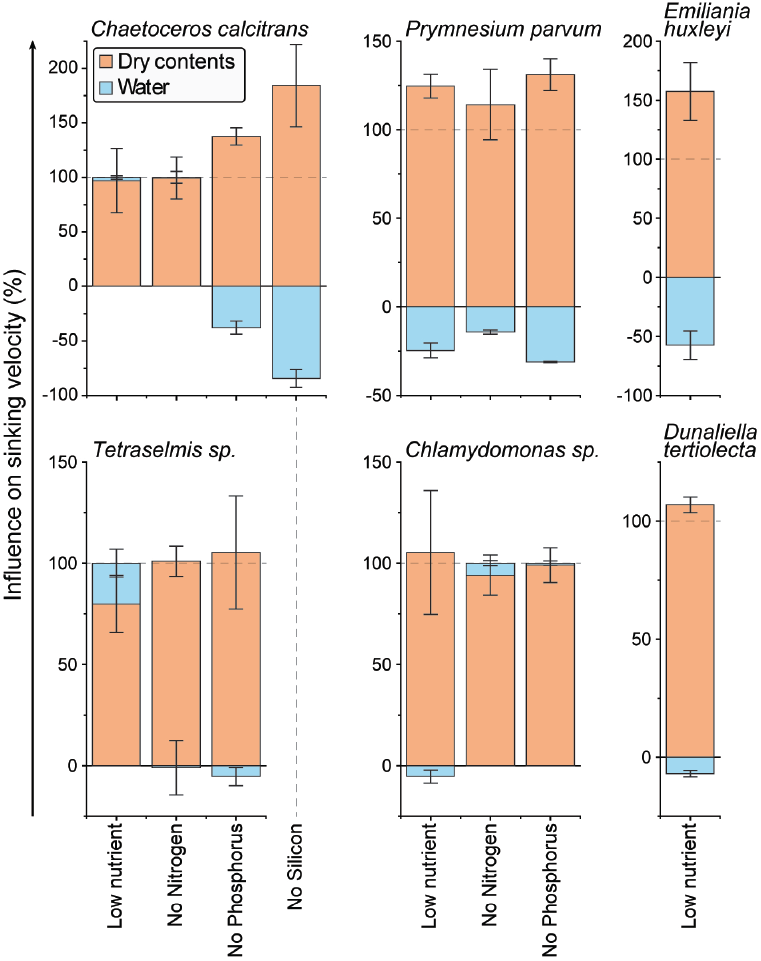
Starvation-induced cell sinking is driven by dry content accumulation. The relative influence of cellular water (blue) and dry contents (orange) changes on cell sinking velocity changes under each starvation condition. Negative values imply that dry and water content changes have opposite effects on the sinking velocity. Data depicts mean±SEM.

### Increased lipid and decreased protein content explain the decreased cell sinking in starving

#### Phaeodactylum tricornutum

To further understand which macromolecules impact cell sinking following starvation, we examined the cell lipid and protein compositions. We first focused on explaining sinking changes in *Phaeodactylum tricornutum*, the only species in our study that decreased sinking following starvation. We fluorescently labeled neutral lipids, which are the principal form of storage lipids ^37^, and cellular proteins in fixed cells and measured the cells using flow cytometry. Lipid accumulation was significantly larger in *Phaeodactylum tricornutum* than in other species tested (Fig S6A). Starved *Phaeodactylum tricornutum* cells contained ∼10-fold more lipids than cells under high nutrient condition (Fig 5A), consistent with previous reports establishing this species as a high lipid producer ^37,38^. Fluorescence and transmission electron microscopy (TEM) revealed that the neutral lipids formed 1-2 large lipid droplets under starvation, whereas the lipids were distributed into smaller droplets under the high nutrient condition (Figs 5B,C). We did not observe obvious changes in the frustule thickness (Fig 5C). When analyzing cellular protein content, we found that *Phaeodactylum tricornutum* displayed a larger decrease than other species did when starved (Figs S3B, S6B), with protein content decreasing ∼4-fold (Fig 5D). According to our simulations (Fig 1C), the lipid accumulation and protein loss following starvation in *Phaeodactylum tricornutum* are both sufficient to explain the decreased sinking (Fig 5E). However, as the combined effect exceeds the experimentally derived sinking, additional compositional changes must also exist (Fig 5E).

**Figure 5.**
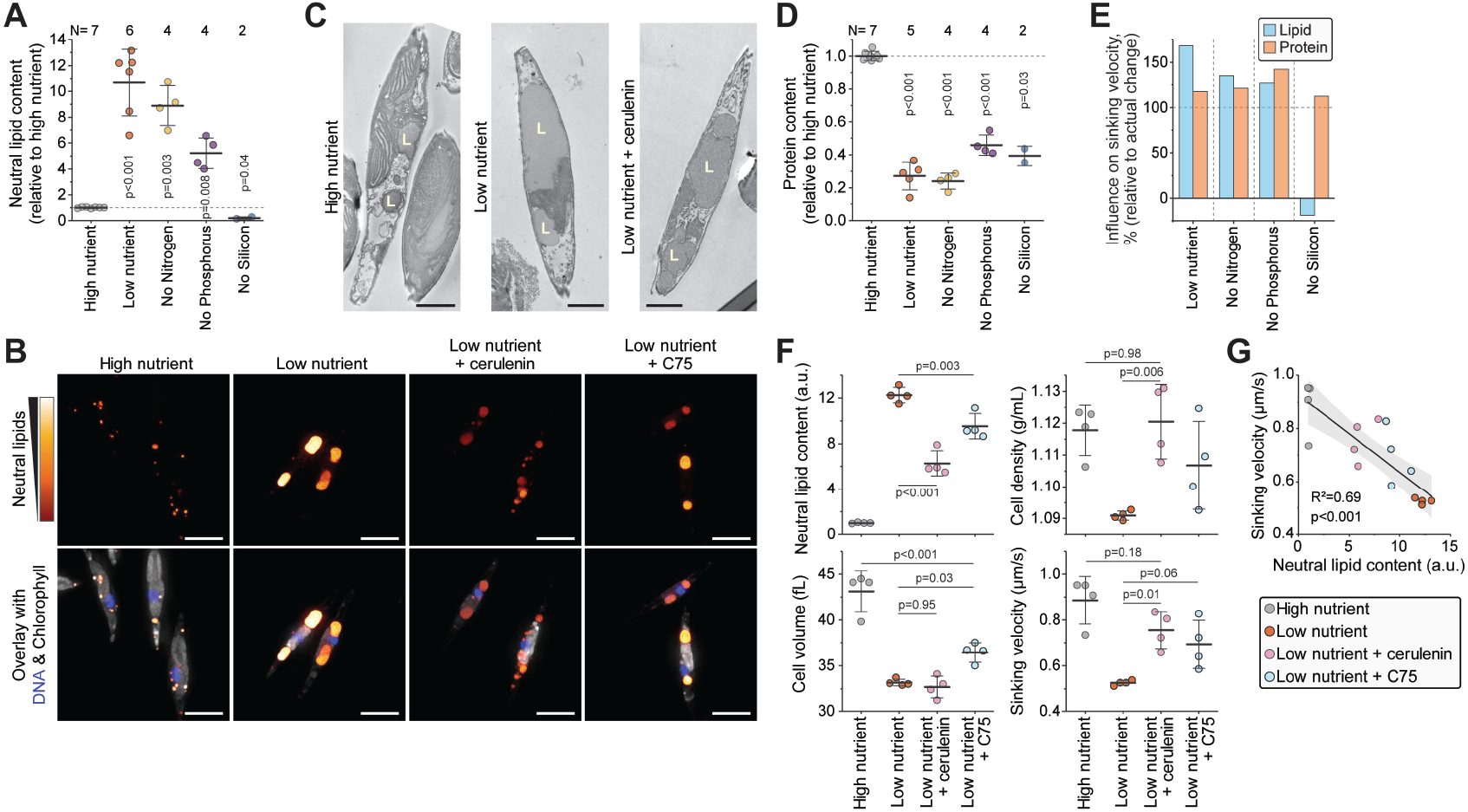
Extensive lipid accumulation and loss of proteins explains the decreased sinking velocity in starving *Phaeodactylum tricornutum*. **(A)** Relative changes in cellular neutral lipid content following 5-day culture under indicated starvation conditions. N depicts the number of independent cultures (dots), bar and whiskers depict mean±SD. **(B)** Fluorescence imaging of neutral lipids (red to yellow), DNA (blue), and chlorophyll (grey) following 5-day culture under indicated conditions. Scalebars depict 5 µm. n>40 cells per condition. **(C)** TEM images following 5-day culture under indicated starvation conditions. Scalebars depict 2 µm. Letter L indicates lipid droplets. n>25 cells per condition. **(D)** Same as (A), but data is for cellular protein content. **(E)** Simulated influence of lipid and protein content changes on cell sinking velocity. Data is normalized to the actual sinking velocity changes observed. **(F)** Neutral lipid content, cell density, volume, and sinking velocity of cells under indicated conditions. Dots depict separate cultures (N=4), bar and whiskers depict mean±SD, p-value obtained by Student’s t-test. **(G)** Correlation between cell neutral lipid content and sinking velocity in the data in (F). Data in all panels is from the species *Phaeodactylum tricornutum*.

We then examined if we could rescue the sinking decrease in *Phaeodactylum tricornutum* by preventing lipid accumulation. We treated cells with two fatty acid synthase inhibitors, cerulenin and C75, for the duration of the low nutrient starvation. We then imaged the cells for neutral lipids (Fig 5B), and quantified neutral lipid content, cell density, volume and sinking (Fig 5F). Cerulenin partly reversed the lipid accumulation and lipid droplet morphology observed under low nutrient starvation, as was cell sinking velocity (Figs 5B,C,F). Cerulenin fully rescued the cell density decrease observed under low nutrient state, but it did not rescue the cell volume decrease. C75 treatment yielded more modest rescues of lipid content and cell sinking than the cerulenin treatment. Overall, the neutral lipid content and sinking of the cells treated with and without fatty acid synthase inhibitors were correlated (p<0.001, ANOVA, R^2^=0.69) (Fig 5G), indicating that a majority of the low nutrient starvation driven sinking can be attributed to lipid accumulation.

Other species, including *Prymnesium parvum* and *Chlamydomonas sp.*, also increased cellular lipid content when starved (Fig S6A), despite increased cell density (Figs 3A, S3A). The lipid accumulation in *Chlamydomonas sp.* was localized exclusively to the cell periphery, where lipid droplets protruded against the plasma membrane (Fig S7A-B). This cellular organization is different from the widely studied freshwater counterpart *Chlamydomonas reinhardtii* ^39^. These lipid droplets increased 2-fold in diameter, but not in number, upon starvation (Figs S7C-D). In addition, *Chlamydomonas sp.* decreased its protein content by ∼2-fold when starved. These lipid and protein content changes are expected to decrease *Chlamydomonas sp.* sinking. As we did not observe this experimentally, we expect additional compositional changes to counteract the lipid increase and protein loss in *Chlamydomonas sp*.

### Increased starch reservoirs can explain the increased cell sinking in starving green algae

Green algae are capable of accumulating both starch and lipid reservoirs under starvation, as carbon fluxes are directed from biosynthesis to storage molecules ^28,29^. However, the accumulation of starch and lipids would have opposing effects on cell sinking (Fig 1C).

This motivated us to examine starch accumulation in green algae. We carried out TEM imaging of the *Tetraselmis sp*., *Chlamydomonas sp*., and *Dunaliella tertiolecta* (Figs 6A, S8A). We observed extensive starch granule accumulation in all three species when starved, with up to ∼40% of the cell area being occupied by starch in *Dunaliella tertiolecta* (Figs 6B). This starvation-induced increase in starch content was due to both the number and size of the starch granules increasing (Figs S8B-C). In addition, we observed that phosphorus starved *Chlamydomonas sp*. cells accumulated less starch than nitrogen starved cells, although this difference was not observed in *Tetraselmis sp.* (Fig 6B). Instead, in *Chlamydomonas sp*., phosphorus starvation resulted in the appearance of acidocalcisome-like organelles ^40^, which may also contribute to sinking (Figs S8A,D).

**Figure 6.**
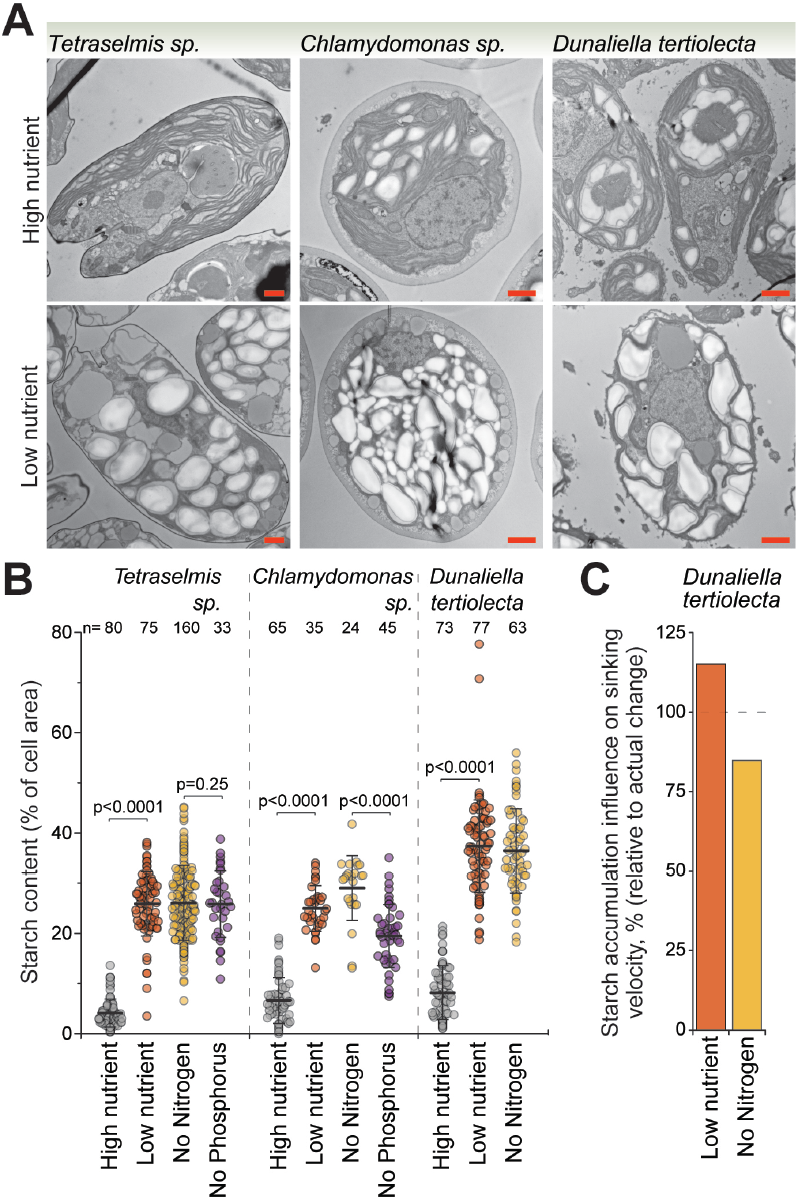
Extensive starch accumulation can explain sinking velocity increases in starving green algae. **(A)** TEM imaging of green algae species following 5-day culture under indicated starvation conditions. Scalebars depict 1 µm. **(B)** Quantifications of cellular starch granule content from TEM images. Dots depict separate cells, bar and whiskers depict mean±SD, n depicts the number of cells measured, and p-values were obtained using ANOVA followed by Tukey’s posthoc test. **(C)** Simulated influence of starch accumulation on cell sinking velocity in *Dunaliella tertiolecta*. Data is normalized to the actual sinking velocity changes observed.

We used *Dunaliella tertiolecta* as a model system for simulations of the influence of starch accumulation on sinking. We chose this species because it displayed little lipid or fractional water content changes when starved (Figs S5, S6), suggesting that sinking may rely more exclusively on carbohydrate accumulation. According to our simulations, the accumulation of starch alone explains approximately all the sinking increase observed in low nutrient starved and nitrogen starved *Dunaliella tertiolecta* (115% and 85%, respectively) (Fig 6C). More broadly, across all three green algae species, the starch accumulation had a larger simulated impact on sinking than changes in cell lipid, protein, or water content (Fig S8E). Thus, our results indicate starch accumulation as an important molecular mechanism responsible for green algae sinking under nutrient starvation.

## DISCUSSION

Our study revealed that 8 out of 9 tested phytoplankton species alter their sinking when starved for nutrients. In most tested species, this can be explained by the accumulation (or loss) of dry contents, and we have exemplified how the accumulation of lipids and carbohydrates, and the loss of proteins, can function as mechanisms to alter cell sinking. However, it is important to recognize that additional mechanisms that influence sinking can also exist. Some phytoplankton can accumulate large amounts of pigment molecules ^30^, polyphosphate storages ^40^, or inorganic components, such as silica and calcium carbonate, all of which may increase cell sinking. Curiously, our experiments did not reveal any conditions where phytoplankton become buoyant in seawater. As shown by our simulations, this would require a large accumulation of water in the absence of other dry content accumulation, or a very large accumulation of lipids. Both mechanisms are likely to require extraordinary changes to intracellular organization, as shown in *Pyrocystis noctiluca* ^31^, and may therefore be rare. Alternatively, phytoplankton would have to lose >70% of their dry contents, which would likely compromise many cellular functions, or accumulate extremely low-density contents, such as gas vesicles, which are currently only reported to exist in prokaryotes ^41,42^. Thus, while our work illustrates several mechanisms used to increase eukaryotic phytoplankton sinking, mechanisms that achieve buoyancy are still largely unknown.

An unexpected discovery in our study is that most phytoplankton grow larger when starved. While the increased cell size is easily explained by decreased proliferation rates ^43^, it is in a stark contrast to most model systems, where cell size decreases with nutrient starvation ^33,44,45^. Why would phytoplankton have evolved to increase their size upon starvation? Increased cell sinking is one possible explanation, but not the only one. If phytoplankton sink significantly deeper in the water column, they will be exposed to less light, and an increase in cell size (area) could enable more light harvesting. Additionally, increased cell size can store more energy (lipids and carbohydrates), which may support cell viability and motility in the light-limited deeper waters. A better understanding of the size-dependence of metabolic processes within a species will help elucidate this ^46^.

We also find that most phytoplankton are denser when starved than when proliferating, as observed in several other model systems from bacteria to humans ^33,47–49^. This suggests that there may exist a more fundamental “starvation state” where cellular properties are adjusted to cope with starvation, possibly to conserve energy ^48^. We did not observe systematic cell size increases under starvation, suggesting that this starvation state is distinct from that observed upon genome dilution, where cells enter starvation-like state due to excessive cell size increases in the absence of DNA replication ^50,51^. Importantly, we also identified a few exemptions to this behavior, such as the starvation of *Emiliania huxleyi*, where density did not increase. Future studies comparing these differential starvation responses may elucidate the physiological consequences of the high-density starvation state.

The changes in cell sinking that we observe following starvation do not necessarily reflect regulation of cell sinking, as they could be byproducts of starvation-induced metabolic effects. However, if sinking is regulated (so that it promotes cell fitness), our results can provide context for such regulation. The gravitational sinking velocity of most species studied here is under 1 m/day, which makes long distance (*e.g.*, 50 m) vertical migration very slow and unlikely, especially for small cells. However, in a competitive environment, a fitness advantage might be gained by much more modest changes in depth. This is especially true for cells that inhabit depths close to the nutricline. In addition, in some species, gravitational sinking could be augmented by motility, and cells may also acquire higher sinking velocities by aggregating together. Although our studies are technically limited to species which do not display excessive aggregation, our results nonetheless suggest that the increased cell sinking velocity following starvation could promote cell-to-cell encounters. However, we note that the *in situ* vertical migration of phytoplankton is under significant influence by turbulence and ocean currents, and further investigation is required to understand the importance of gravitational sinking to the movement of phytoplankton.

Finally, our results could support modeling of ocean ecosystems and nutrient cycles. Our study has connected the macromolecular content of cells to their sinking, and the macromolecular content is also indicative of the C:N:P ratio of cells ^30,52^. It seems therefore likely that cells, as well as cell aggregates, with different elemental ratios sink at different rates, which could contribute to marine carbon and nutrient cycles. Linking elemental stoichiometry and macromolecular content to the vertical movement of cells and cell aggregates is an important area of future modeling efforts.

## MATERIALS AND METHODS

### Simulations of cell sinking velocity

The gravitational sinking velocity of a cell is a function of cell size, density, and shape, as described in *Eq (1)*. All phytoplankton species in this study can be modelled as spheroid or ellipsoid (Fig S3), with a multiplicative shape correction factor less than 10% compared to a sphere ^53^. Thus, cell shape was not considered in the simulation, and all cells were assumed to be spherical.

We modelled a cell as the collection of all its intracellular molecules and categorized these molecules into five groups: proteins, lipids, carbohydrates, water, and others. Hence, the cell volume is the sum volume of individual molecules, *V* = Σ *V*_*i*_, and the cell density is the weighted average of the densities of those molecules (Table S1), *ρ* = Σ *w*_*i*_· *ρ*_*i*_, where the subscript *i* refers to each molecule group and *w*_*i*_ refers to their volume fraction. Once the cell volume and density are defined by the molecular composition, the sinking velocity can be calculated using *Eq (1)* for any given cell state.

Using our model, we simulated sinking velocities of *Phaeodactylum tricornutum, Dunaliella tertiolecta, Chaetoceros calcitrans*, and a hypothetical average species. We first determined their molecular compositions under high nutrient conditions from literature values (Table S2). For each species, we then varied the volume of one molecule group at a time or, in the case of ‘dry contents’, all non-water molecule groups together, while the other parameters remained constant.

### Influence of biophysical properties and intracellular molecules on sinking velocity

To determine the influence of biophysical properties (i.e. cell volume and density) on cell sinking, we used a first order Taylor expansion near the baseline condition (*ρ*_0_, *r*_0_) that represents high-nutrient condition of the cell.

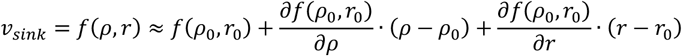

Hence, the influence of cell density and volume, *I*_*density*_ and *I*_*volume*_, respectively, can be defined as the fractions of the two derivative terms.

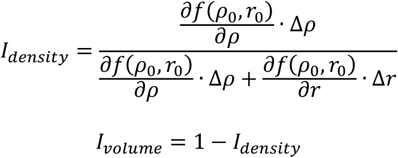

Similarly, the influence of molecular composition on sinking velocity was decoupled into those of water and dry content (proteins, lipids, carbohydrates, and others). In this case, the sinking velocity was rewritten as a function of water volume, dry volume, and dry density (taking water density as a constant).

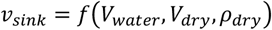

The first order Taylor expansion near the high-nutrient condition 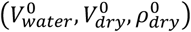 indicates the influences of water volume, dry volume, and dry density. The influence of each variable, *I*_*water*_, *I*_*dry volume*_, or *I*_*dry density*_, is the ratio of each derivative term to the total of all three terms. For example, *I*_*water*_can be rewritten as:

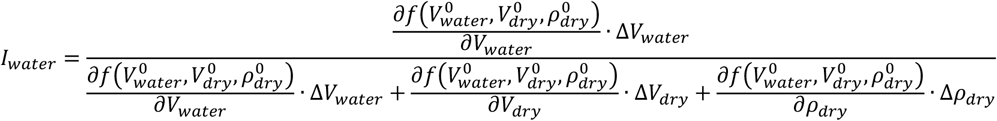

The influence of dry content can be further calculated as the sum of dry volume and dry density as:

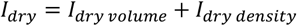

### Phytoplankton species, culture conditions, and drug treatments

All phytoplankton species were obtained from Provasoli-Guillard National Center for Marine Algae and Microbiota (NCMA) and the species belong to the Culture Collection of Marine Phytoplankton (CCMP). All species identifications were verified using light microscopy, as well as TEM for a subset of species. A list of species, their CCMP identifiers, and their maintenance growth media are shown in Table S3.

All algae were cultured as previously ^32^. High nutrient condition corresponds to L1 (or L1-Si) media, low nutrient condition corresponds to the high nutrient conditions with the nutrients being diluted 100-fold, and specific nutrient starvations correspond to the high nutrient condition in the absence of the indicated nutrient. For experiments, maintenance cultures were split 1:4 for 2 days to achieve exponentially growing cultures. These cultures were then split to each indicated nutrient condition and culture for 5 days prior to measurements of cell sinking. The lighting was available from a nearby window at approximately 50 μmol/s/m^2^ of PAR and day length varying between 9 h and 14 h. All cultures were grown at 22°C ± 1.5°C. Key experiments were repeated using a controlled lighting setup with 100 μmol/s/m^2^ of PAR with day length set at 12 h and temperature at 22°C.

Lipid synthesis was inhibited with cerulenin (Cayman Chemical, Cat#10005647) and C75 (Cayman Chemical, Cat#10005270). Lipid synthesis inhibitors were used at 10 µM concentration and the treatment duration corresponded to the low nutrient starvation (5 days).

### Cell buoyant mass and volume measurements

Cell buoyant mass and volume measurements were carried out identically to previous report ^32^. In short, buoyant masses were measured using the SMR ^54^, where a vibrating cantilever with a microfluidic channel detects changes in resonant frequency due to the presence of a cell, which correlates to its buoyant mass. Two SMRs were used: one with an 8 × 8 µm cross-section for smaller phytoplankton cells and one with a 15 × 20 µm cross-section for larger cells. Cell volumes were measured using a Coulter counter (Multisizer 4), based on electric impedance changes as cells passed through an orifice. Two aperture sizes (20 µm and 100 µm) were used depending on cell size. Both SMR and Coulter counter provide measurements of single cells within 1–100 ms per cell, and the measurements were carried out at room temperature in L1-Si/100 media. The measurements were calibrated using NIST-certified polystyrene beads (2–10 µm, Duke Standards, Thermo Scientific). All measurements were carried out between 9:30am and 3:30pm, and paired samples (*e.g.*, high nutrient and corresponding low nutrient sample) were measured within 1 hour of each other.

### Determining cell sinking velocities and Péclet numbers

Phytoplankton gravitational sinking velocities, *v*_*sink*_, were calculated according to (*Eq 1*). The following values were used for environmental constants: the dynamic viscosity of seawater of 1.07 × 10^−3^ Pa·s, and density of seawater of 1.026 g/ml. Population average cell radius was calculated from the cell volume measurements and population average cell density was calculated by comparing the volume and buoyant mass measurements. All phytoplankton species in this study were estimated to have low Reynolds numbers (<10^−6^). We note that the exact gravitational sinking velocities would be influenced by changes in seawater viscosity and density, which could occur if, for example, seawater temperature changed significantly.

Péclet numbers were calculated according to *Pe* = *UL/D*, where *U* is the sinking velocity, *U* is the cell length, and *D* is the external particle’s diffusivity ^55^. We used the diffusivity value of 1,700 μm^2^/s for nitrate and 10 μm^2^/s for a small particle (*e.g.*, viral particle).

### Cell proliferation rate measurements

Cell proliferation rates were derived from Coulter counter -based cell count measurements. Cell counts were measured on three consecutive days, where the middle day corresponded to the cell mass and volume measurements. Proliferation rates were calculated assuming exponential growth.

### Fluorescent labeling of proteins, lipids, and DNA

For all fluorescent labeling approaches, the cells were fixed for 10 min in L1-Si media (or corresponding nutrient limited media) containing 4% formaldehyde, after which the cells were washed twice with PBS. Neutral lipids were stained using 2 µM Bodiby 493/503 (4,4-Difluoro-1,3,5,7,8-Pentamethyl-4-Bora-3a,4a-Diaza-s-Indacene, ThermoFisher, Cat#D3922) in PBS for 20 min. After staining, the cells were washed two times with PBS. Cellular proteins were stained using the amine-reactive LIVE/DEAD Fixable Red Dead Cell Stain (ThermoFisher, Cat#L34972) using 2x supplier recommended concentration in PBS for 15 min ^56^. After staining, the cells were washed with PBS solution containing 5% BSA, and again with PBS. DNA was stained using 10 µg/ml Hoechst 33342 (ThermoFisher, Cat#H3570) in PBS for 20 min. After staining, the cells were washed two times with PBS.

### Fluorescence and brightfield microscopy

Fluorescence and brightfield microscopy samples were plated on poly-L-lysine (Sigma-Aldrich, Cat#P8920) coated glass bottom CELLVIEW plates (Greiner Bio-One) for >30 min prior to imaging. The samples were imaged at RT using DeltaVision wide-field deconvolution microscope with standard DAPI, FITC, TRICT, Cy5, and POL filters, and a 100x oil-immersion objective. z-layers were typically collected with 0.3 µm spacing covering >8 μm in height. Fluorescence image deconvolution was carried out using DeltaVision software.

### Flow cytometry

The flow cytometry samples were identical in preparation to those used for fluorescence microscopy. Instead of plating on imaging plates, the samples were cleaned from aggregates using a 100 μm filter. The samples were then analyzed using BD Biosciences FACS Celesta with 405 nm, 488 nm and 561 nm excitation lasers and 450/40 nm, 515/20 nm, 610/20 nm, and 710/50 nm emission filters. Samples were gated on FSC and SSC to exclude too small particles, and a typical analysis measured 10,000 cells within the FSC/SSC gate. Chlorophyll autofluorescence was used to ensure that all measured particles were phytoplankton cells.

### Water and dry content measurements

Cellular water and dry content, including the density and volume of cell’s dry contents, were determined as detailed before ^32^. In short, the average buoyant mass of cells in a given population was measured with the SMR, as detailed above. These measurements were then repeated after moving the cells to L1-Si/100 media where 90% of the water was deuterated (D_2_O). D_2_O mixes freely with the water inside the cell and D_2_O has a higher density than H_2_O, resulting in a different population average buoyant mass. By comparing these two buoyant mass averages, along with the measurement solution densities (L1-Si_H2O_ and L1-Si_D2O_), we can solve for the average dry volume and the density of the dry volume in the cell population ^57,58^. These measurements were then compared to the Coulter counter -based total cell volume measurements to derive the volume of water inside the cells.

### Electron microscopy

For TEM sample preparation, the cells were fixed for 60 min in L1-Si media (or corresponding nutrient limited media) containing glutaraldehyde and paraformaldehyde at final concentrations of 2.5% and 2%, respectively. After fixation, the cells were washed two times with 100mM sodium cacodylate (pH 7.2). After washing, samples were fixed for 60 min on ice with 1% osmium tetroxide in a solution containing 1.25% potassium ferrocyanide and 100 mM sodium cacodylate. Next, the samples were washed multiple times with 100 mM sodium cacodylate and with 50 mM sodium maleate. The samples were then stained with 2% uranyl acetate in sodium maleate buffer at RT o/n. The samples were rinsed with DI water and dehydrated using a series of 10 min ethanol incubations. The ethanol concentrations varied in ascending order, from 30% to 100%. The samples were then washed twice in propylene oxide for 30 min, and moved to propylene oxide - resin mixture (1:1) for o/n. The samples were moved to a new propylene oxide - resin mixture (1:2) for 6 hours and then into pure resin o/n. The samples were then moved to molds and polymerized at 60°C for 48 hours to form blocks suitable for sectioning. Thin sections of 60 nm were obtained using a Leica UC7 ultramicrotome. The sections were collected on carbon-coated nitrocellulose film copper grids.

TEM imaging was carried out using FEI Tecnai T12 transmission electron microscope and an AMT XR16 CCD camera. Typical imaging was carried out with 120 kV voltage. Imaging magnification varied between experiments. All TEM images shown in the manuscript are representative examples.

For SEM sample preparation, the cells were plated on poly-L-lysine coated cover slides for 40 min prior to fixation. The fixatives, 2.5% glutaraldehyde and 2% paraformaldehyde, were added to the cells on the cover slides for 60 min. The samples were then washed two times with 100mM sodium cacodylate (pH 7.2) and post-fixed for 30 min at +4°C using 1% osmium tetroxide in sodium cacodylate buffer. The samples were rinsed with DI water and dehydrated using a series of 10 min ethanol incubations. The ethanol concentrations varied in ascending order from 35% to 100%. Next, the samples were incubated with ethanol/tetramethyl silane mixtures (50/50 mixture, and 20/80 mixture, respectively) for 15 min each. Then, the samples were washed twice with 100% tetramethyl silane. The wash solution was removed, and the samples were left to dry o/n. The dried samples were sputter coated with gold.

SEM imaging was carried out using Zeiss Crossbeam 540 scanning electron microscope. Typical imaging was carried out with 4 kV accelerating voltage, 600 pA probe current, a working distance of 8 mm, and a magnification of 4000x. All SEM images shown in the manuscript are representative examples.

### Image analysis

Lipid droplets, starch granules, and acidocalcisomes were visually identified from TEM images based on existing literature ^37,59,60^. For quantifications, cells, along with intracellular components of interest, were segmented and quantified using MATLAB (R2023a). For each cell, the cell boundary, lipid droplets, starch granules, and acidocalcisomes were manually defined by a freehand line profile drawn using MATLAB’s Image Processing and Computer Vision toolbox. Intracellular composition was then estimated using the area of a component of interest relative to the total cell area. Script can be found at https://github.com/alicerlam/algae.

### Statistics

In all figures, N refers to the number of independent cultures, n refers to the number of separate cells measured. Particles too small to be viable cells were removed from all final analyses. Statistical tests are detailed in figure legends and all p-values were calculated using OriginPro (2025) software.

## Data availability

All cell sinking velocity, buoyant mass, volume, density, proliferation rate, water volume, dry volume, and dry density, data are available in Table S4.

## Declaration of interests

S.R.M. is a co-founder of and affiliated with the companies Travera and Affinity Biosensors, which develop techniques relevant to the research presented.

## Author contributions

Y.W. carried out simulations. Y.W., V.K.K., T.R.U., J.L., and T.P.M. carried out phytoplankton culture and live cell measurements. V.K.K., M.B., A.K.R.L-J., and T.P.M. carried out cell imaging. R.A., A.Z.L., A.R.L., and T.P.M., carried out image analysis. S.R.M. and T.P.M. provided resources and supervised the work. Y.W. and T.P.M. conceived and designed the research and wrote the manuscript with input from all authors.

## Funding and additional acknowledgements

This work was supported by the National Science Foundation (Award 2319028) to S.R.M. The work was also supported, in part, by the Koch Institute Support(core) grant P30-CA14051 from the National Cancer Institute. We thank the Koch Institute’s Robert A. Swanson (1969) Biotechnology Center, specifically the Peterson (1957) Nanotechnology Materials Core Facility (RRID: SCR_018674), the Microscopy Core Facility, and the Flow Cytometry Core Facility for their support. We also thank Prof A. Babbin for useful feedback.

## Notes

### Summary of Updates

Main text has been updated, with most changes in the introduction and discussion sections. Supplemental data file has been updated. Figures have not changed.

