## Supplementary Information for "Biophysical and molecular mechanisms responsible for phytoplankton sinking in response to starvation"

**Affiliations:**
[1] Koch Institute for Integrative Cancer Research, Massachusetts Institute of Technology, Cambridge, MA 02139, USA.
[2] Harvard-MIT Department of Health Sciences and Technology, Institute for Medical Engineering and Science, Massachusetts Institute of Technology, Cambridge, MA 02139, USA.
[3] Department of Biological Engineering, Massachusetts Institute of Technology, Cambridge, MA 02139, USA.
[4] Department of Mechanical Engineering, Massachusetts Institute of Technology, Cambridge, MA 02139, USA.

[*] These authors contributed equally to this work


**This file contains:**
Supplementary Figures S1-S8
Supplementary Tables S1-S3
Supplementary References

**Additional supplementary files not included here:**
Supplementary Table S4

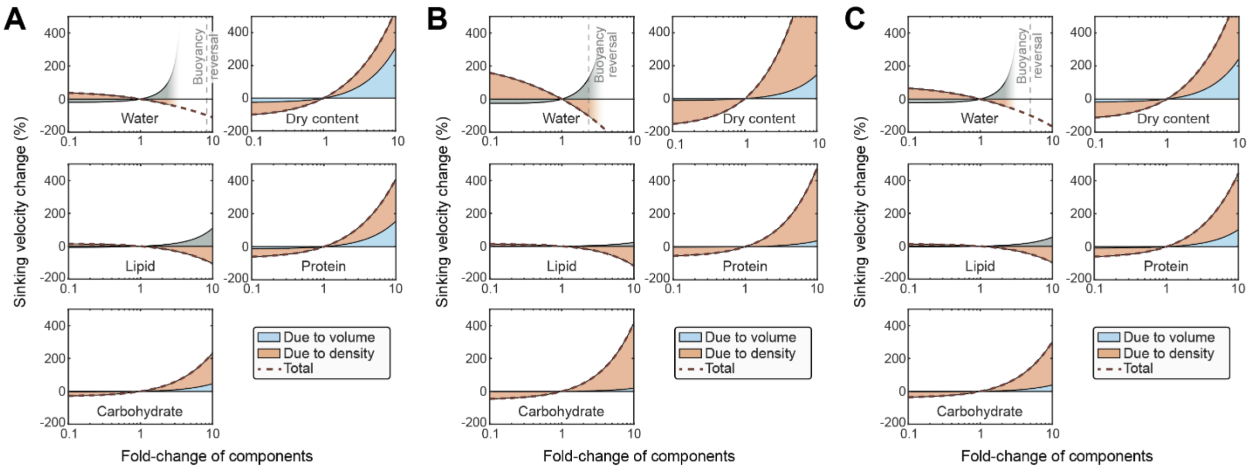


**Figure S1. Simulations of cell sinking are largely independent of starting cell composition.**
**(A-C)** Simulations of cell sinking velocity in a typical green alga (A, *Dunaliella tertiolecta*), diatom (B, *Chaetoceros calcitrans*), and a hypothetical average species (C) as a function of changing cell composition. Sinking velocity changes due to cell volume and cell density changes are separated in blue and orange, respectively, and total cell sinking change is depicted by red dotted line. Changes in dry content refer to corresponding changes in all other contents except water. For large water content increases, separation of volume and density contributions are excluded. See Tables S1-S2 for more details.

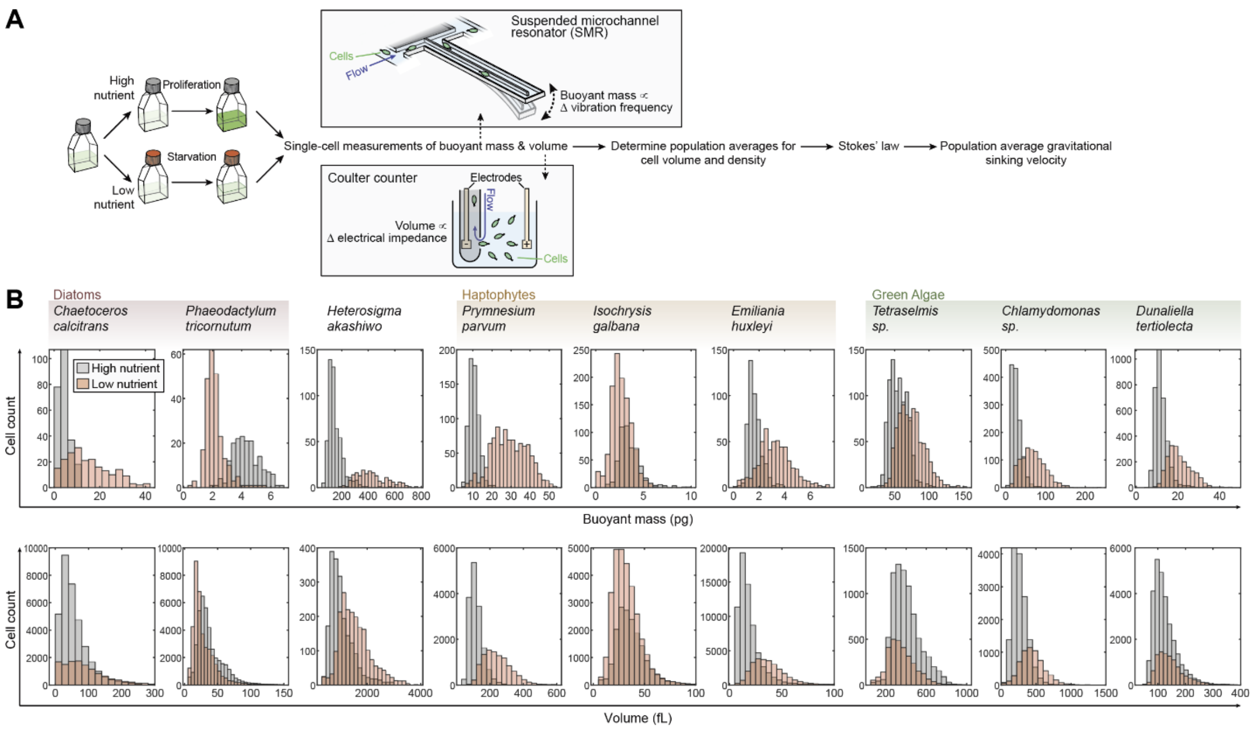


**Figure S2. Single-cell mass and volume responses to starvation.**
**(A)** Schematic of workflow for measuring single-cell buoyant masses and volumes, which were then used to determine population average density, volume, and gravitational sinking velocity. SMR is a microfluidic device, where a cell is flown through a channel embedded in a vibrating cantilever. The change in the cantilever's vibration frequency is proportional to the buoyant mass of the cell. Coulter counter is a fluidic device, where a cell is flown through a small aperture, displacing the measurement solution (artificial seawater). This changes the electrical resistance across the aperture in a manner proportional to cell volume. **(B)** Representative single-cell buoyant mass (top) and volume (bottom) histograms following a 5-day culture under high (grey) and low (orange) nutrient conditions in indicated species.

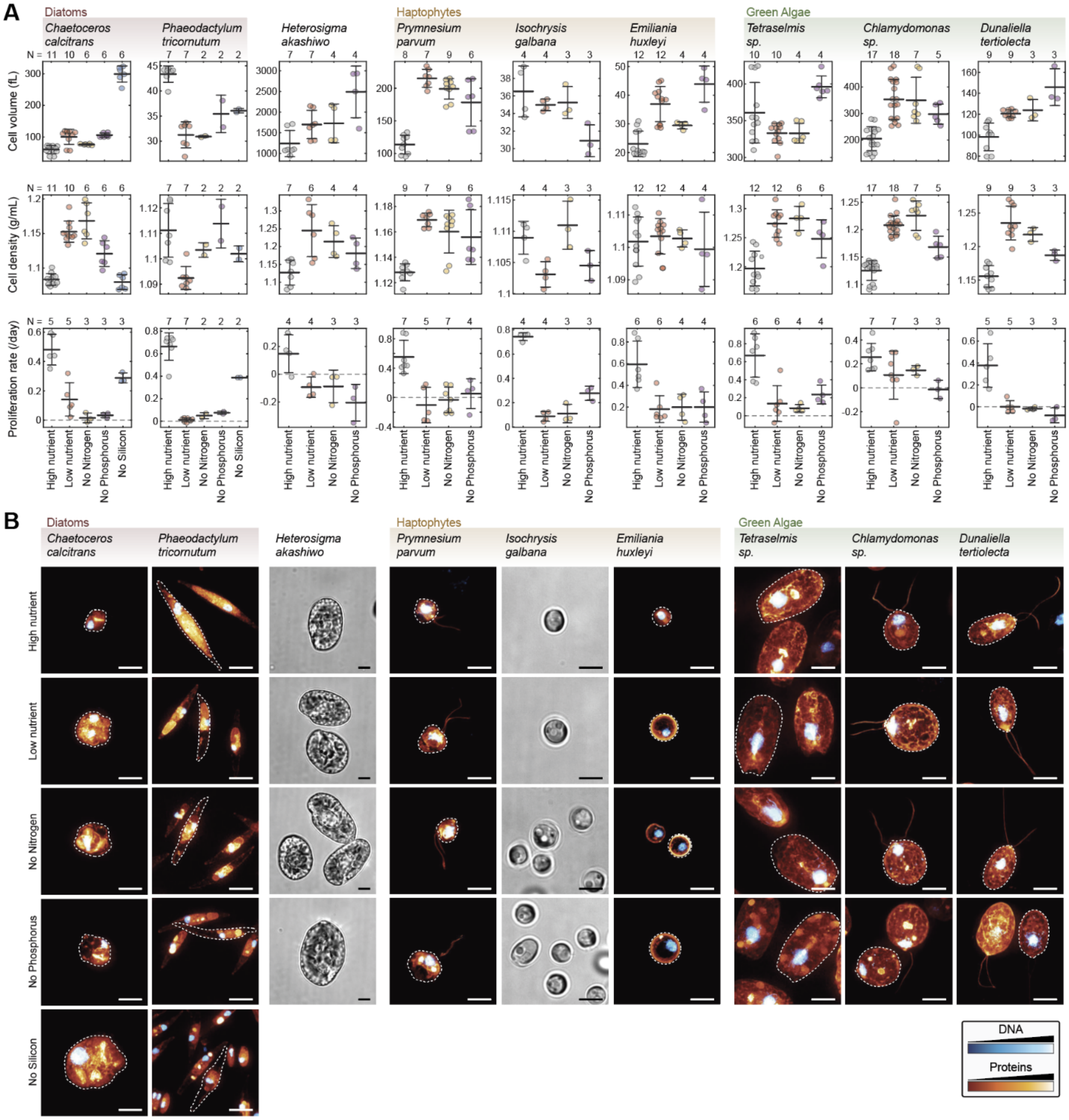


**Figure S3. Phytoplankton cell volumes and densities change when starved, but cell shapes do not.**
**(A)** Cell volume (top row), density (middle row) and proliferation rate (bottom row) under indicated nutrient conditions in indicated species. Dots depict independent cultures, bars and whiskers depict mean±SD, N depicts the number of independent cultures. Note that the *Emiliania huxleyi* strain is non-calcifying. **(B)** Representative fluorescence microscopy images of indicated species following a 5-day culture under indicated nutrient conditions. The cells were labeled for total protein content (red-to-yellow) and DNA content (blue-to-white). Cell outlines are highlighted with a dotted line for one cell in each fluorescence image. Brightfield images were used for instead of fluorescence imaging for *Heterosigma akashiwo* and *Isochrysis galbana*. All scalebars denote 5 µm. Note that *Heterosigma akashiwo* is zoomed out 2x in comparison to other images. n > 20 cells per condition. *Phaeodactylum tricornutum* cells were near-exclusively in the fusiform morphotype.

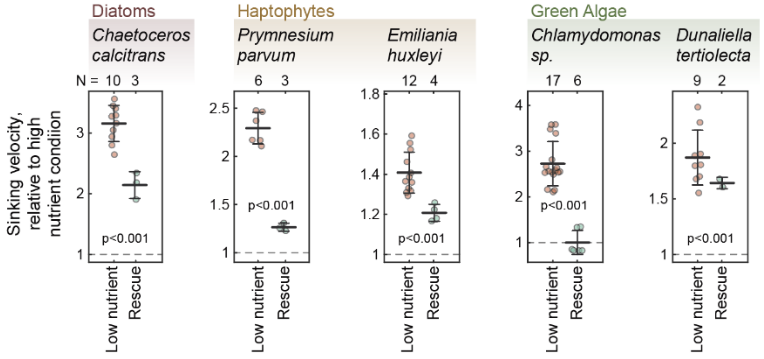


**Figure S4. Starvation-induced cell sinking velocities are reversible.**
Gravitational sinking velocities were measured following 5-day starvation under low nutrient condition after which nutrients were resupplied and sinking velocities were measured again 2 days later (rescue). All data are normalized to high nutrient condition on day 5. p-values depict Welch's t-test between low nutrient and rescue conditions.

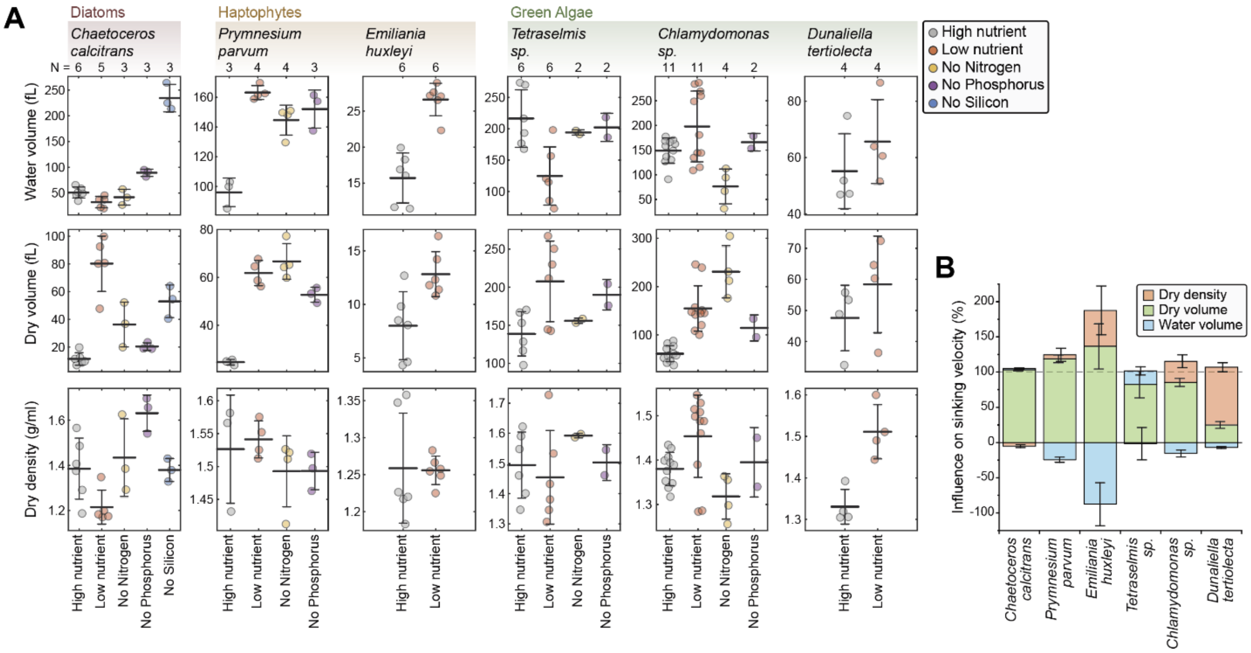


**Figure S5. Cellular dry and water content following starvation.**
**(A)** Volume of water (top row) and dry contents (middle row), and the density of the dry contents (bottom row) in an average cell in indicated species following a 5-day culture under indicated starvation conditions. N depicts the number of independent cultures (dots), bar and whiskers depict mean±SD. **(B)** The relative influence of cellular water volume (blue), dry volume (green), and the density of dry content (orange) changes on cell sinking velocity changes under low nutrient starvation condition. Data depicts mean±SEM.

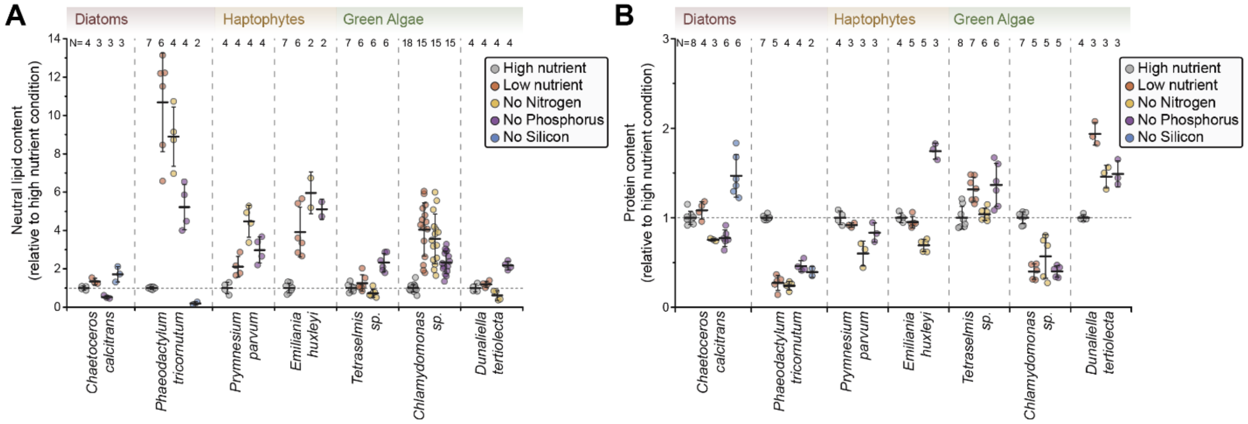


**Figure S6. Lipid and protein content changes following starvation**

**(A)** Relative changes in cellular neutral lipid content following 5-day culture under indicated starvation conditions. Data normalized to high nutrient condition within each species. N depicts the number of independent cultures (dots), bar and whiskers depict mean±SD. **(B)** Same as (A), but data is for cellular protein content. Data for *Phaeodactylum tricornutum* is a repeat of the data shown in Figs 5A, D. See Fig S3B for example fluorescence images of the cellular protein content labeling.

Note that the data represent neutral lipid and protein content per cell, but as many species increase their cell volume following starvation, the concentration of neutral lipids and proteins may differ significantly from the changes visualized here.

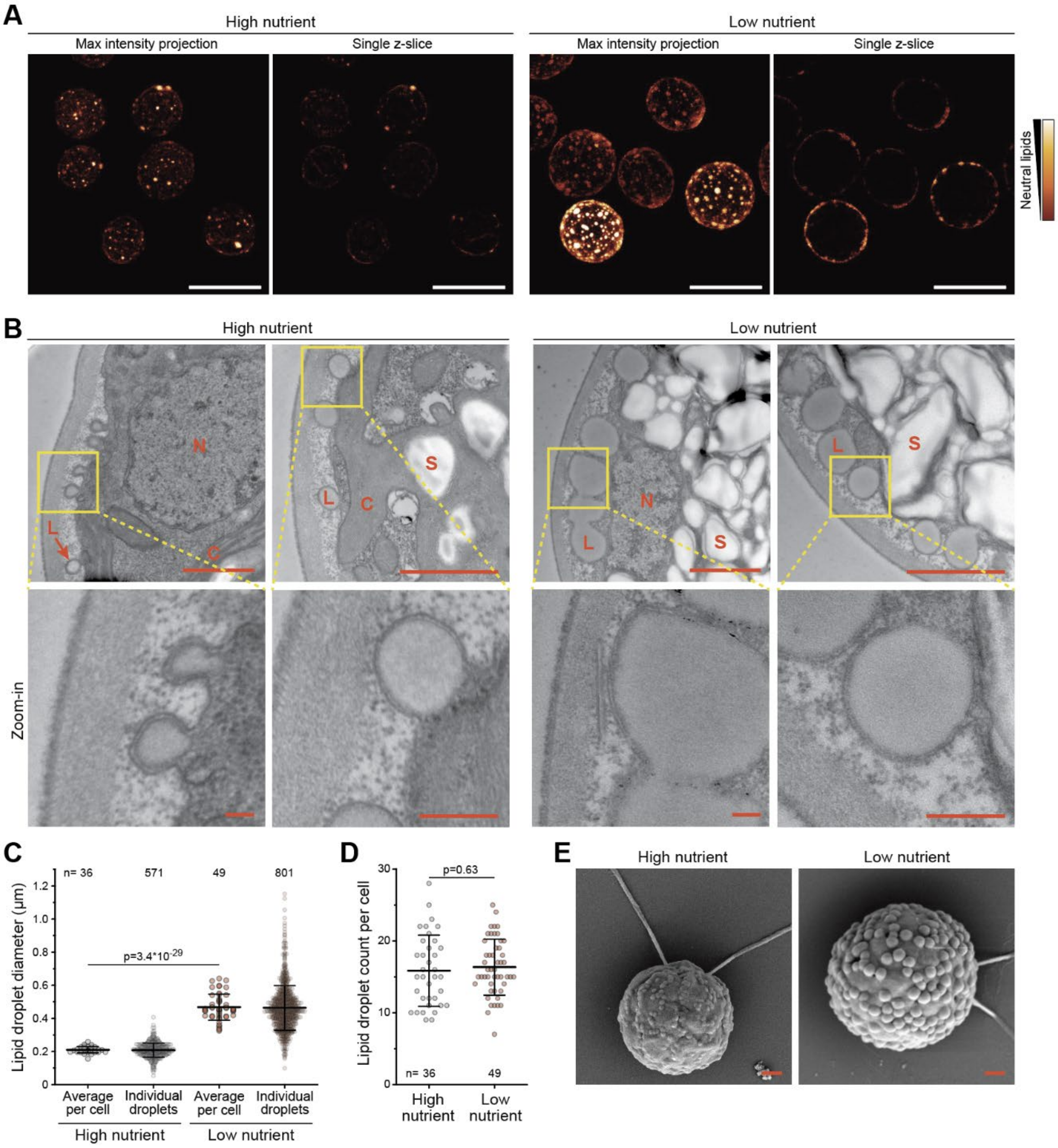


**Figure S7. *Chlamydomonas sp*. accumulates lipid droplets exclusively at the cell periphery.**
**(A)** Fluorescence microscopy of neutral lipids in *Chlamydomonas sp*. cells following 5-day culture under high and low nutrient conditions. Both maximum intensity and single z-layer images are shown to highlight the lipid droplet localization at cell periphery. Scale bars denote 10 µm. **(B)** TEM of *Chlamydomonas sp*. cells following 5-day culture under high and low nutrient conditions. Zoom-ins (bottom row) visualize lipid droplets. Key cell compartments are indicated with letters (C = chloroplast, L = lipid droplet, N = nucleus, S = starch granule). Scale bars denote 1 µm on top row and 200 nm on bottom row (zoom-ins). **(C-D)** Quantifications of lipid droplet diameter (C) and count (D) from TEM images, p-value depicts Welch's t-test. **(E)** Scanning electron microscopy (SEM) of *Chlamydomonas sp*. cells. Cell dehydration and shrinkage during SEM sample prep reveals 'bumps' on the cell surface, which are identical in size and location to the lipid droplets observed in TEM and fluorescence microscopy. Scalebars denote 1 µm.

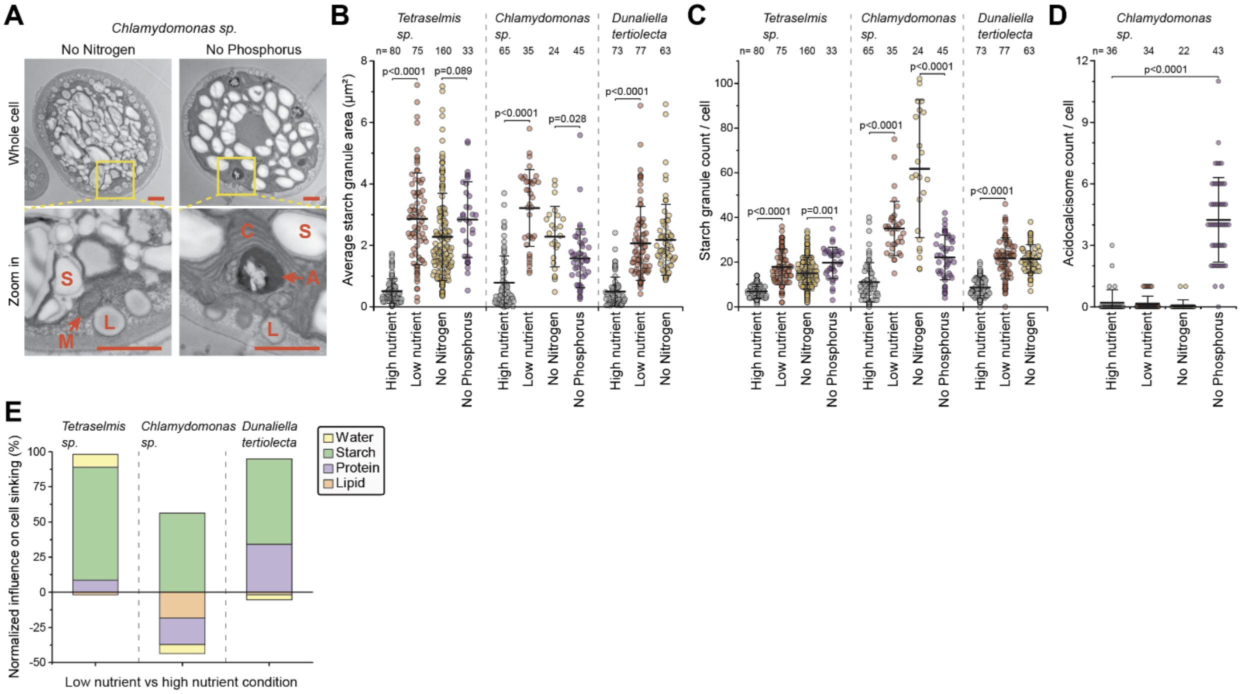


**Figure S8. Starving green algae accumulate larger starch granules and, in some cases, acidocalcisome-like organelles.**

**(A)** TEM imaging of *Chlamydomonas sp.* following 5-day culture under nitrogen and phosphorus starvation. Scalebars depict 1 µm. Key cell compartments are indicated with letters in the zoom in (A = acidocalcisome, C = chloroplast, L = lipid droplet, M = mitochondria, S = starch granule). **(B-C)** Quantifications of average starch granule area (B) and number per cell (C) from TEM images. Dots depict separate cells, n depicts number of cells analyzed, bar and whiskers depict mean±SD. **(D)** Quantifications of acidocalcisome number per cell from TEM images. Acidocalcisomes are electron-dense, round organelles in TEM images, which often store calcium, phosphate, and metals. Analysis was limited to *Chlamydomonas sp.,* as acidocalcisomes were not observed in other green algae. Dots depict separate cells, n depicts number of cells analyzed, bar and whiskers depict mean±SD. **(E)** Simulated influence of water, starch, protein, and lipid content changes on cell sinking velocity in the green algae species. The analysis was limited to comparing high and low nutrient conditions, where cellular content changes were determined most comprehensively. Data is normalized so that the components add up to 100%. Note that for *Chlamydomonas sp.* the starch accumulation can counteract sinking velocity influence of lipid accumulation and protein loss, but additional mechanism(s) must be present to fully explain the experimentally determined cell sinking.

In panels B-D, p-values were obtained using ANOVA followed by Tukey's posthoc test.

**Table S1. Molecular (dry) density values used in simulations.**

| *Molecular species* | *Density (g/ml)* | *Reference* | *Additional notes* |
|---|---|---|---|
| *Protein* | 1.35 | [1,2] (BNIDs 110540, 114284) | Generic protein values range from 1.31 to 1.37 |
| *Lipid* | 0.92 | [1] (BNID 114321) | Combined values of triolein, trilinolein, and tricaprylin |
| *Carbohydrate* | 1.5 | [1] (BNIDs 103206, 112354, 112354) | Combined values of starch granules |
| *Water* | 1.0 | | Reflects $H_2O$ independently of any salts |
| *Other* | 1.3 | | Estimated density of all other cellular components |

**Table S2. Cell composition values used in simulations.** The fractional (w/w) dry content values are derived from literature and the fractional water content values are derived from experiments in this paper.

| *Species name* | *Dry content* | | | | *Water* | *Ref.* |
|---|---|---|---|---|---|---|
| | ***Protein*** **(w/w dry)** | ***Lipid*** **(w/w dry)** | ***Carbohydrate*** **(w/w dry)** | ***Other*** **(w/w dry)** | **(v/v total)** | |
| *Dunaliella tertiolecta* | 49% | 17% | 17% | 18% | 52.6% | [3–5] |
| *Chaetoceros calcitrans* | 29% | 13% | 18% | 40% | 80.9% | [6,7] |
| *Phaeodactylum tricornutum* | 45% | 9% | 18% | 29% | 67.1% | [8,9] |
| *Generic* | 45% | 15% | 20% | 20% | 66.7% | |

**Table S3. Details of the phytoplankton species studied.**

| *Species name* | *CCMP identifier* | *Culture media* | *Motility* | *Additional notes* |
|---|---|---|---|---|
| *Chlamydomonas sp.* | CCMP222 | L1-Si | Motile | |
| *Dunaliella tertiolecta* | CCMP362 | L1-Si | Motile | |
| *Phaeodactylum tricornutum* | CCMP632 | L1 | Non-motile | |
| *Prymnesium parvum* | CCMP708 | L1-Si | Motile | Toxic |
| *Tetraselmis sp.* | CCMP908 | L1-Si | Motile | |
| *Chaetoceros calcitrans* | CCMP1315 | L1 | Non-motile | |
| *Isochrysis galbana* | CCMP1323 | L1-Si | Motile | |
| *Heterosigma akashiwo* | CCMP1870 | L1-Si | Motile | Toxic |
| *Emiliania huxleyi* | CCMP2090 | L1-Si | Non-motile | Non-calcifying |
